# Distinct roles for the Anterior Temporal Lobe and Angular Gyrus in the spatio-temporal cortical semantic network

**DOI:** 10.1101/544114

**Authors:** Seyedeh-Rezvan Farahibozorg, Richard N. Henson, Anna M. Woollams, Olaf Hauk

**Affiliations:** MRC Cognition and Brain Sciences Unit, University of Cambridge, Cambridge, UK, CB2 7EF; Wellcome Centre for Integrative Neuroimaging, Nuffield Department of Clinical Neurosciences, University of Oxford, Oxford, UK, OX3 9DU; Neuroscience and Aphasia Research Unit, School of Biological Sciences, University of Manchester, Manchester, UK, M13 9PL

**Author notes:** Corresponding author: Seyedeh-Rezvan Farahibozorg.

## Abstract

It is now well recognised that human semantic knowledge is supported by a large neural network distributed over multiple brain regions, but the dynamic organisation of this network remains unknown. Some studies have proposed that a central semantic hub coordinates this network. We explored the possibility of different types of semantic hubs; namely “representational hubs”, whose neural *activity* is modulated by semantic variables, and “connectivity hubs”, whose *connectivity* to distributed areas is modulated by semantic variables. We utilised the spatio-temporal resolution of source-estimated Electro-/Magnetoencephalography data in a word-concreteness task (17 participants, 12 female) in order to: (i) find representational hubs at different timepoints based on semantic modulation of evoked brain activity in source space; (ii) identify connectivity hubs among left Anterior Temporal Lobe (ATL), Angular Gyrus (AG), Middle Temporal Gyrus and Inferior Frontal Gyrus based on their functional connectivity to the whole cortex, in particular sensory-motor-limbic systems; and (iii) explicitly compare network models with and without an intermediate hub linking sensory input to other candidate hub regions using Dynamic Causal Modelling (DCM) of evoked responses. ATL’s activity was modulated as early as 150ms post-stimulus, while both ATL and AG showed modulations of functional connectivity with sensory-motor-limbic areas from 150-450ms. DCM favoured models with one intermediate hub, namely ATL in an early time window and AG in a later time-window. Our results support ATL as a single representational hub with an early onset, but suggest that both ATL and AG function as connectivity hubs depending on the stage of semantic processing.

## 1 Introduction

How are the networks supporting conceptual knowledge organised in the brain? Previous literature has focussed on two brain subnetworks, one comprising heteromodal areas in Perisylvian cortex that support all types of conceptual knowledge, and one comprising sensory-motor-limbic areas that are only recruited for certain categories of concepts (Binder & Desai 2011). However, the precise role of these areas and their inter-connectivity remain unknown (Pulvermüller 2013). Most previous studies have attempted to localise semantics in the brain using neuropsychology or functional magnetic resonance imaging (fMRI) (Binder *et al.* 2009; Lambon Ralph *et al.* 2016). Nonetheless, these methodologies do not provide the temporal resolution required to separate retrieval of semantic information from post-retrieval processes such as mental imagery (Hauk 2016; Pulvermüller *et al.* 2009) or to uncover fast changes in the communication between different nodes within the semantic network.

We used Electro-and Magneto-EncephaloGraphy (EEG/MEG) to compare dynamic brain activity and connectivity in response to visually-presented concrete and abstract words. We used this contrast to examine the role of hetermodoal semantic areas (hereafter referred to as candidate semantic “hubs”), considering that these two categories differ with respect to their general semantic processing demands and general sensory-motor-affective attributes (Binder *et al.* 2005; Dhond *et al.* 2007). The term hub is often used to refer to the functional role of brain areas in semantic brain networks (Patterson *et al.* 2007; Pulvermüller 2013). However, graph theory has provided a number of empirical metrics to characterise hubs (Bullmore & Sporns 2009). Here, we considered two types of heteromodal hubs that are linked to neuroscientific theories of semantics: “representational hubs”, in which the activity of neural populations is modulated by the semantic variables, and “connectivity hubs”, which mediate communication between semantic areas through modulations of connections to heteromodal and sensory-motor-limbic nodes of the semantic network. This distinction is related to the distinction between “representation” and “connectivity” made by Woollams and Patterson (2017) for describing the role of the anterior temporal lobe (ATL) within a “hub-and-spokes” model.

The hub-and-spokes model, based on computational modelling, neuropsychological research on semantic dementia patients (Patterson *et al.* 2007; Snowden *et al.* 2017), and neuroimaging research on healthy participants (Lambon Ralph *et al.* 2016; Lau *et al.* 2013), proposes that a single semantic hub binds together brain regions within a distributed semantic network, and underlies category-general semantics (Lambon Ralph *et al.* 2016; Rogers *et al.* 2004). However, meta-analytic neuroimaging evidence also implicates several other brain regions (Binder *et al.* 2009), including posterior inferior parietal lobe (especially angular gyrus, AG), middle temporal gyrus (MTG) and inferior frontal gyrus (IFG), as potential hubs. Indeed, neuro-computational modelling of word learning promotes the possibility of multiple semantic hubs (Tomasello *et al.* 2017). Some authors have therefore argued for an “interactive continuum of hierarchically ordered neural ensembles” (Binder & Desai 2011), similar to the framework of convergence zones (Barsalou 2009; Martin 2016; Meyer & Damasio 2009).

Here, we used a largely data-driven approach, in order to objectively characterise the brain networks at different stages of semantic processing. First, we analysed evoked activity to find the brain areas that first distinguished concrete and abstract words, revealing possible candidates for representational hubs. Second, we examined functional connectivity between the main hub candidates (ATL, IFG, MTG, AG) and distributed semantic areas, particularly sensory-motor-limbic systems, in order to identify connectivity hubs in the whole-cortex networks. Third, using Dynamic Causal Modelling (DCM) of evoked responses (David *et al.* 2006), we tested for the presence of a central connectivity hub within the heteromodal semantic subnetwork that links sensory input to different nodes of this network in an early and a later time-window. For this purpose, we constructed a hierarchy of model comparisons comprising two levels, and asked: a) are models with a single connectivity hub preferred over models with no hubs, and b) in the preferred models, do the areas that function as connectivity hubs change from earlier to later stages of semantic retrieval?

## 2 Materials and methods

### 2.1 Data acquisition and pre-processing

#### 2.1.1 Participants

20 healthy native English speakers participated in the study, but 3 participants were removed due to excessive movement artefacts or measurement error. Hence 17 participants (age 27±6 years, 12 female) entered the final analysis. A mean handedness laterality quotient of 82 (min 41, max 100) was obtained from a reduced version of the Oldfield handedness inventory (Oldfield 1971). All participants had normal or corrected-to-normal vision with no reported history of neurological disorders or dyslexia. The experiment was approved by the Cambridge Psychology Research Ethics Committee and volunteers were paid for their time and effort.

#### 2.1.2 Stimuli

Participants were presented with 184 monomorphemic abstract and concrete words (92 per category, see Extended Data Table 1 - 1 for the list of words), matched for a number of psycholinguistic variables including Kucera-Francis (KF) and CELEX word frequencies, familiarity, concreteness and imageability ratings as well as the number of letters/phonemes/syllables (see Table 1 for details). KF Frequency, Familiarity, Concreteness and Imageability values were taken from the MRC Psycholinguistic Database (Coltheart 1981) and CELEX Frequency taken from the MCWord Database (Binder & Medler, 2005). The two categories differed significantly on concreteness and imageability (ts>19.3575, ps<.0005) as indicated by independent samples t-tests, but not with respect to the other aforementioned variables ((ts<1.28, ps>.202).

**Table 1.** **Psycholinguistic properties of stimuli:** average and standard deviations for relevant psycholinguistic variables for the semantic decision stimuli as a function of concreteness. List of stimuli are presented in Table 1 - 1 (Extended Data).

| | Concrete (average ± std) | Abstract (average ± std) |
| --- | --- | --- |
| **Number of Letters** | 5.9 ± 0.8 | 6.0 ± 0.8 |
| **Number of Phonemes** | 5.0 ± 1.2 | 5.2 ± 1.2 |
| **Number of Syllables** | 2.0 ± 0.6 | 2.1 ± 0.8 |
| **KF Frequency** | 15.0 ± 18.0 | 15.0 ± 15.0 |
| **CELEX Frequency** | 12.0 ± 15.0 | 15.0 ± 14.0 |
| **Familiarity Rating** | 462.0 ± 59.0 | 447.0 ± 98.0 |
| **Concreteness Rating** | 590.0 ± 26.0 | 303.0 ± 39.0 |
| **Imageability Rating** | 566.0 ± 41.0 | 388.0 ± 78.0 |

KF: Kucera-Francis; std: standard deviation.

#### 2.1.3 Procedure and task

Single-word stimuli appeared as 28-point Arial font in white on a black screen within a visual angle of 4 degrees in a slightly dimmed and acoustically shielded MEG chamber at the MRC Cognition and Brain Sciences Unit, University of Cambridge. Duration of stimulus presentation was 150 ms, with an average SOA of 2400 ms (uniformly jittered between 2150 and 2650 ms). Participants preformed a concreteness decision task, by making button presses with their right hand, using index and middle finger to distinguish concrete and abstract words. Short breaks were included after about every 50 trials. Participants were given a few minutes of practice prior to the experiment, using different stimuli, until they felt comfortable with the task. The first two trials (filler items) after each break and at the beginning of each block were not included in the analysis.

A concreteness judgement task was preferred over a passive task in order to ensure that participants stay alert throughout the experiment. Additionally, the contrast of concrete and abstract words has been extensively used in the previous literature in order to identify the heteromodal semantic areas (Binder 2016; Binder *et al.* 2005; Dhond *et al.* 2007) based on modulations of brain activity. Here, we extended the previous explorations by incorporating the temporal trajectory of brain activity and whole-cortex statistics in order to identify the representational hubs in the whole-cortex. In addition, this contrast was considered suitable for identifying connectivity hubs through modulations of connections among the heteromodal semantic areas as well as connectivity between heteromodal and sensory-motor cortices. More specifically, concrete words are generally more strongly associated with sensory-motor attributes while abstract words have been suggested to rely more on the heteromodal subnetwork of semantics (Binder 2016). Thus, this contrast was considered more suitable to tackle representational and connectivity hubs compared to more general (e.g. word/pseudoword) or more specific (fine-grained categories of concepts) semantic contrasts.

#### 2.1.4 EEG/MEG data acquisition and pre-processing

MEG data were acquired in a magnetically shielded room using a Neuromag Vectorview system (Elekta AB, Stockholm, Sweden), with 204 planar gradiometers and 102 magnetometers (i.e. 306 channels overall). EEG data were collected concurrently using a 70-electrode EEG cap (EasyCap GmbH, Herrsching, Germany). EEG reference and ground electrodes were attached to the nose and left cheek, respectively. The Electro-Oculo-Gram (EOG) was recorded by placing electrodes above and below the left eye (vertical EOG) and at the outer canthi (horizontal EOG). Data were acquired with a sampling rate of 1000Hz and a band pass filter of 0.03 to 330 Hz. Prior to the MEG recording, the positions of 5 Head Position Indicator (HPI) coils attached to the EEG cap, 3 anatomical landmark points (two ears and nose) as well as approximately 50-100 additional points covering the whole EEG cap were digitised using a 3Space Isotrak II System (Polhemus, Colchester, Vermont, USA) for later co-registration with MRI data. It is worth noting that simultaneous EEG/MEG recordings have been shown to improve the accuracy of source localisation which motivated their concurrent recording in the current study (Molins *et al.* 2008; Sharon *et al.* 2007).

Our analysis pipeline for the data is illustrated in Figure 1. The first step of data pre-processing included applying signal-space separation (SSS) implemented in the Maxfilter software (Version 2.0) of Elekta Neuromag to the raw MEG data in order to remove noise from sources distant to the sensor array (Samu Taulu and Matti Kajola 2005). The Maxfilter software also involved movement compensation and bad channel interpolation for MEG data. All the next analysis steps (except Dynamic Causal Modelling) were performed in the MNE-Python software package (http://martinos.org/mne/stable/index.html) (Gramfort *et al.* 2013, 2014). Raw data were visually inspected for each participant, and consistently bad EEG channels were marked and interpolated. Data were then FIR band-pass filtered between 1 and 48 Hz with a window length of 40s in both forward and backward directions to achieve zero phase delay. Independent Component Analysis (ICA) was applied to the filtered data in order to remove eye movement and heart artefacts. We used FastICA algorithm (Hyvärinen *et al.* 2000) as included in scikit-learn python package (Pedregosa *et al.* 2011) and implemented in MNE-Python meeg-preprocessing package (with minor manual changes to achieve a better artefact rejection for some participants). After ICA, data were divided into epochs from −500ms to 700ms around the word onsets. Epochs were rejected if peak-to-peak amplitudes were higher than the following thresholds, based on previous norms: 120 µV in the EEG (except for 2 cases where we increased the threshold to 150 µV, because high rejection rates could be identified as due to excessive Alpha activity despite good behavioural performance), 2500 fT in magnetometers, 1000 fT/cm for gradiometers. Trials with incorrect responses were also excluded from further analysis.

**Figure 1.**
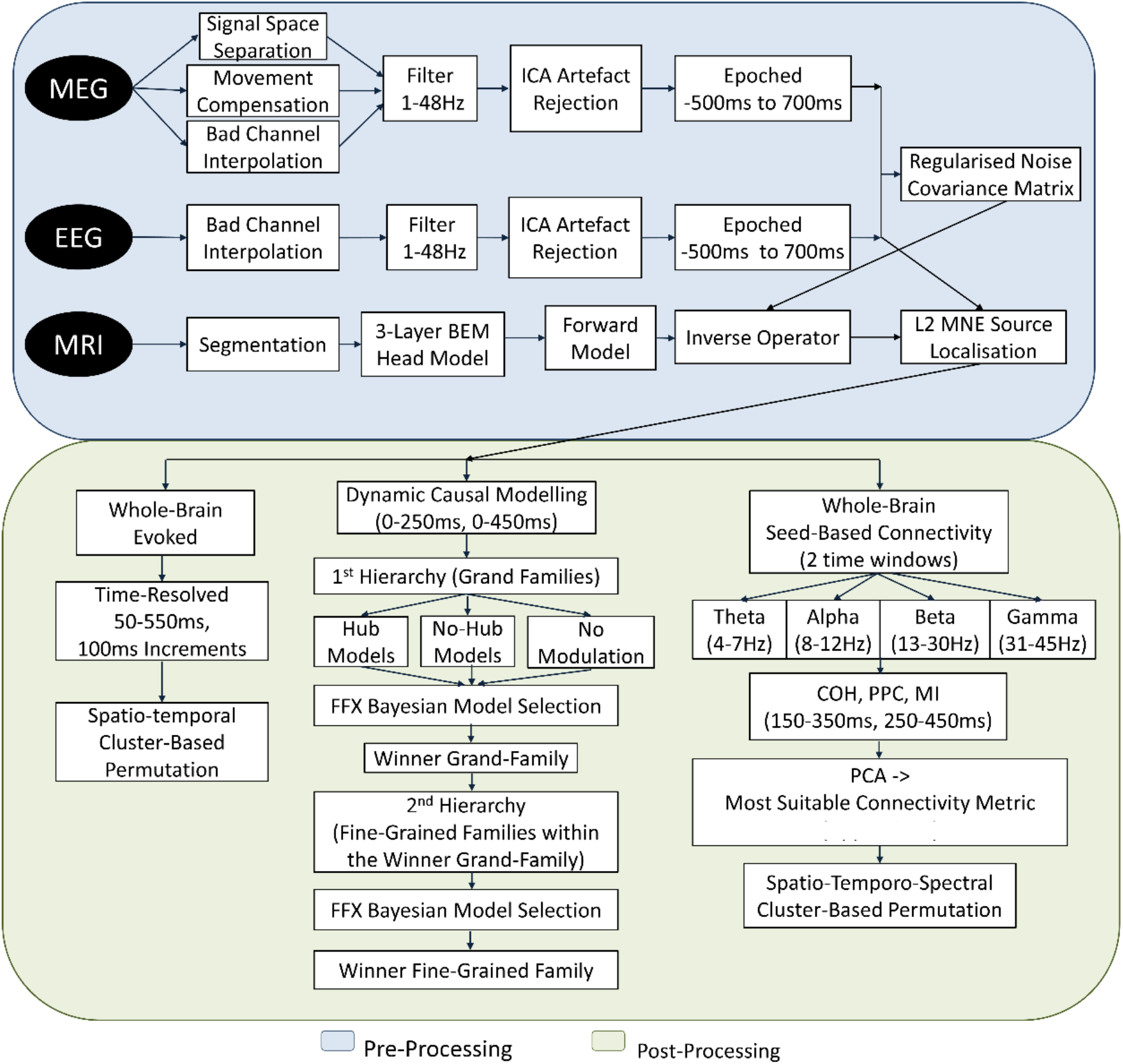
A flowchart of different steps of data analysis.

#### 2.1.5 Forward model and inverse solution

We used MNE-Python software to compute forward and inverse models. The forward model was computed based on a Boundary Element Model (BEM) of the head derived from structural MR images for each participant. EEG/MEG sensor configurations and MRI images were co-registered based on the aforementioned digitisation points. Structural MRI images were processed using the automated segmentation algorithms in FreeSurfer software (Version 5.3; http://surfer.nmr.mgh.harvard.edu/) in order to obtain the reconstructed scalp surface (Dale *et al.* 1999; Fischl *et al.* 1999). The result of the FreeSurfer segmentation was processed further using MNE software package (Version 2.7.3) and the original triangulated cortical surface, which included more than 160,000 vertices per hemisphere, was down-sampled to a tessellated grid in which the average edge of each triangle was approximately 2.5mm (Segonne *et al.* 2004). A three-layer BEM containing 5120 triangles per layer was created for EEG and MEG from scalp, outer skull surface and inner skull surface, respectively. The noise covariance matrices for each dataset were computed and regularised in a single framework which computes the covariance using a diagonal regularisation technique with regularisation factor of 0.1 for all the channel types. Baseline intervals of 500ms duration pre-stimulus were used for noise covariance estimation. The resulting regularised noise covariance matrix was used to assemble the inverse operator for each participant using L2 minimum-norm estimation (L2 MNE) with a loose orientation constraint value of 0.2 and without depth weighting.

### 2.2 Whole-cortex evoked analysis

This analysis was aimed at identifying representational hubs in the whole cortex. After removing bad trials according to aforementioned criteria, the number of epochs were equalised between concrete and abstract words by matching the time of trial presentation, i.e, removing excessive epochs for concrete words that showed minimal temporal alignment with the abstract words. Equalisation of the number of trials was performed so as to remove any potential biases due to the differences in signal to noise ratios (SNR), given that error rates were significantly higher for the abstract words (section 3.1). Trials for each condition were averaged in sensor space in order to yield an evoked response per participant and condition, which were then projected onto the source space using L2 MNE. We used MNE-Python’s default SNR = 3.0 for regularisation of the inverse operator for evoked responses. Afterwards, the individual participant results were morphed to the standard average brain (fsaverage5) in Freesurfer software, yielding time courses of activity for 20,484 vertices for each participant and condition. Source-estimated time courses were then averaged in five time windows from 50ms to 550ms with 100ms increments for further analysis.

### 2.3 Seed-based connectivity analysis

We computed whole-cortex seed-based connectivity with the main left-hemispheric heteromodal semantic areas as seeds: ATL, MTG, AG and IFG (see Figure 6) in order to identify connectivity hubs based on the modulation of their connections to distributed semantic areas. Meta-analytic evidence suggests the heteromodal subnetwork of semantics to be bilateral but stronger in the left hemisphere (Binder et al., 2009), particularly for verbal stimuli (Rice *et al.* 2015). Therefore, we constrained the seed regions to the left hemisphere but measured their connectivity to all other brain vertices. For this purpose, we extracted the ROI time courses and computed Magnitude Square Coherence (COH, chosen following a procedure described in 2.3.1) between each ROI time course and every vertex in the brain in four frequency bands of Theta (4-7 Hz), Alpha (8-12 Hz), Beta (13-30 Hz) and Gamma (31-45 Hz) (Engel & Fries 2010) and in early (150-350ms) and late (250-450ms) time windows post-stimulus. Note that the time windows for connectivity analysis have the same onsets as those used in 2.2 except that (a) we used longer time windows here (200ms vs. 100ms) in order to ensure a reliable spectral functional connectivity estimation and (b) we restricted this analysis to only two time windows, because connectivity analysis requires corrections for multiple comparisons across vertices, times, frequency bands and seeds (see 2.4 for details). The onset of these time windows (i.e. 150 and 250ms) was informed by the earliest semantic effects reported in previous literature (Hauk 2016; Moseley *et al.* 2013). In order to extract each ROI time course, we first identified a vertex within the ROI that showed the highest sensitivity to that ROI. To this aim, we computed cross-talk functions (CTFs) and identified the vertex inside that ROI that showed the largest CTF value (for details of ROI_CTF_ calculations refer to (Farahibozorg *et al.* 2018; Hauk *et al.* 2011; Liu *et al.* 1998)). Thereafter, we extracted ROI time courses based on this vertex for each epoch. Next, in order to compute COH, the phase and amplitude/phase consistency were calculated using a multitaper algorithm using the default implementation in MNE-Python (version 0.9). Note that different approaches have been used in the previous literature in order to summarise ROI time courses, including averaging voxel time courses, eigenvectors and using voxels with maximum power (Colclough *et al.* 2015). Considering the limitations of the spatial resolution of the EEG/MEG source localisation, we preferred voxels that showed maximum sensitivity to an ROI’s signal (i.e. most likely to receive signal from each specific ROI).

Magnitude-squared Coherence (COH) is the absolute value of the complex-valued Coherency, which describes the degree of covariance between the amplitudes and phases of two signals (Nunez *et al.* 1997). Coherency is formulated as:

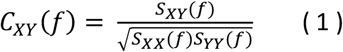

where X(f) and Y(f) are Fourier transforms of the brain signals, f denotes frequency, S_XY_ is the cross-spectrum and S_xx_ and S_yy_ denote auto-spectra of X(f) and Y(f). It is worth noting that S is Fourier transform of the cross-covariance, and hence coherency can be thought of as the analogue of cross correlation in the frequency domain. C_ij_(f) is complex with real and imaginary parts which contain amplitude and phase. Magnitude squared Coherence between two signals is measured as the absolute value of C_ij_(f) in Equation 1.

#### 2.3.1 Choice of Coherence as the most-suitable connectivity metric

COH was chosen among several alternatives in a data-driven manner (details can be found in Extended Data Figure 5 - 1 and Table 5 - 1). Due to the complexity of the neuronal dynamics measured by EEG/MEG, numerous methods have been proposed to quantify connectivity between cortical sources, each designed to capture one or a few aspects of the signals (e.g. Bastos & Schoffelen 2016; Greenblatt *et al.* 2012). Therefore, selecting one particular connectivity method for a specific purpose would require a detailed knowledge of the neuronal mechanisms of brain connectivity. However, such mechanisms for semantic networks have not yet been established. Here, we identified the most suitable method for our data utilising a novel approach based on Principal Component Analysis (PCA) (Jung *et al.* 2000; Lagerlund *et al.* 1997). PCA provides a robust multivariate method of dimensionality reduction and feature selection that has been used frequently at different stages of EEG/MEG pre-/post-processing (Jung *et al.* 2000; Lagerlund *et al.* 1997). By projecting the multidimensional data on orthogonal axes (aka. PCs), PCA can find similarities and differences between the connectivity estimations yielded by different metrics and project them on a single PC and distinct PCs, respectively. Additionally, it can identify principal axes along which the maximum variance of a data is explained as well as the original connectivity metrics that are highly correlated to the most prominent (i.e. first) PC. These metrics can be regarded as the most suitable connectivity metrics (MSC) for a data.

We started by sub-selecting a few connectivity metrics theoretically by focusing on the key methods of functional connectivity in three families: phase plus amplitude coupling, phase coupling and information theoretic (focused on probability distributions). Next, we selected one method from each family, to address the questions of this study: magnitude-squared Coherence (COH), Pairwise Phase Consistency (PPC) and mutual information (MI) (Greenblatt *et al.* 2012). Considering that these three metrics might measure similar and/or different aspects of the data, we utilised PCA in order to: 1) find similarities between them; 2) find unique aspects to each method; 3) identify the method with the highest correlation to the first PC as the most suitable connectivity metric.

After estimating seed-based connectivity using each of the three metrics, for each condition, we concatenated the whole brain connectivity vectors (length N_vertices_) from all participants, times, seeds, frequency bands and connectivity methods in one matrix yielding grand connectivity matrices CMc and CMa of size N_vertex_ × N_participants_ × N_seeds_ × N_times_ × N_bands_ × N_conn_ for concrete and abstract words, respectively. We concatenated CMc and CMa alongside the first dimension and obtained a CM matrix. Thereafter, PCA was computed on 2D sub-matrices obtained from the first and last dimension of CM for each participant, time, seed and band (i.e. sub-matrix size 2N_vert_ × N_conn_) with dimension reduction along the second dimension (i.e. connectivity methods).

We computed variance explained by each PC as well as correlation of each PC to each connectivity method. The explained variance and correlation values were then averaged over times, seeds and participants in order to yield one value for each connectivity method at each frequency band. In order to account for the potential non-normalities or differences in the probability distribution of different connectivity methods, Box-Cox transform (Sakia 1992) was used to obtain a Gaussian distribution for each connectivity method before conducting PCA. Furthermore, considering the fact that COH, PPC and MI have different scales, we conducted weighted PCA (i.e. normalised by variance across rows of each 2N_vert_xN_conn_ sub-matrix) on centred data.

We found that COH and PPC were highly correlated with the first PC (correlation 0.85 or higher) which explained more than 50% of the variance for every frequency band. MI was partially correlated with the spectral measures; however, it was predominately projected on the second PC that explained approximately 30% of the variance of the data. Based on these findings, COH and PPC were identified as the more suitable measures for the data. Considering that the former is sensitive to both amplitude and phase couplings while the latter is only sensitive to the phase, we selected COH as the representative connectivity metric for the rest of the study.

#### 2.3.2 Choice of seeds

Candidate semantic hubs in ATL, IFG, MTG and AG were defined based on the key heteromodal semantic areas proposed in the previous literature, in particular with reference to the meta-analytic evidence by (Binder *et al.* 2009) and a more recent review by (Pulvermüller 2013). It is worth noting that subdivisions of the temporal cortex in heteromodal semantics are not fully established, and previous studies have considered different number of subregions (Binder *et al.* 2009; Jackson *et al.* 2018; Lambon Ralph *et al.* 2016; Pulvermüller 2013). Here, we defined MTG based on the aforementioned meta-analytic evidence, except informed by studies of semantic dementia and the hub-and-spokes model of semantics (Lambon Ralph *et al.* 2016; Patterson *et al.* 2007), we defined the anterior part of the temporal lobe as an independent ROI. Additionally, we defined Angular Gyrus and parts of the Supramarginal Gyrus (SMG) as one seed labelled “AG”. This seed has been identified as a key semantic area by (Binder *et al.* 2009).

### 2.4 Whole-cortex statistical analysis: cluster-based permutation

We used a cluster-based permutation test (Maris & Oostenveld 2007) for statistical analysis of the whole-cortex evoked and seed-based functional connectivity results. For this purpose, we computed univariate vertex-wise t-tests and thresholded them at a t-value corresponding to an initial p-value p0 (two-tailed). Cluster-based permutation was applied to these thresholded t-maps and randomisation was replicated 5000 times in order to obtain the largest random clusters. The cluster-level significance for the original clusters was then calculated as the percentile of the cluster size compared to the largest random clusters across the 5000 permutations. We used spatio-temporal clustering for the whole-cortex evoked responses (accounting for multiple comparisons across vertices and time windows) and spatio-temporo-spectral clustering for the seed-based connectivity in order to also take the four frequency bands into account. Furthermore, considering that cluster-based permutation results can be sensitive to the choice of p0 (Smith & Nichols 2009), we tested several initial thresholds and only clusters that appeared based on more than one p0 were deemed robust. More specifically, for the whole-cortex evoked responses, we tested five thresholds (0.05, 0.045, 0.04, 0.025 and 0.01) and for the seed-based connectivity we tested stricter thresholds (0.025, 0.01, 0.008, 0.002 and 0.001) to make some allowance for the multiple comparisons across the four seeds (i.e. candidate hubs). Additionally, considering the low spatial resolution of EEG/MEG of our source localisation method for deeper brain areas (see (Farahibozorg *et al.* 2018; Hauk *et al.* 2011; Liu *et al.* 1998) for more details), before conducting cluster-based permutation, the areas highlighted in green in Figure 2 were excluded.

**Figure 2.**
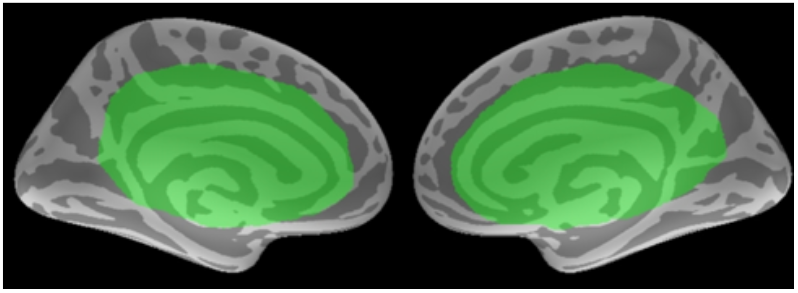
**Vertices excluded from whole-cortex statistical analyses:** green labels defined manually (informed by the previous studies; e.g. (Farahibozorg *et al.* 2018; Hauk *et al.* 2011; Liu *et al.* 1998)) mark deeper brain areas that were removed from the whole-cortex statistical analysis due to the limited spatial resolution of the EEG/MEG source localisation.

### 2.5 Dynamic Causal Modelling (DCM)

DCM analysis focused on identification of the organisation of connectivity among the aforementioned candidate hubs: left ATL, IFG, MTG and AG, as well as the visual word form area (vWFA) in the posterior fusiform gyrus of the left hemisphere which was used as the input region. As the first step, we computed evoked source estimates in the same manner as outlined in 2.2 with two exceptions. Firstly, since DCM for ERP requires signed evoked responses (i.e. reflecting the direction of current flow), we here computed source reconstructed ERPs for dipole components perpendicular to the cortical surface based on the aforementioned source estimates with loose orientation constraint. Secondly, in order to obtain more compatibility with the previous DCM ERP literature (Chennu *et al.* 2016; Garrido *et al.* 2008; Phillips *et al.* 2015), we used a band-pass filter between 1-35 Hz. Next, we used CTFs to identify the vertex with the highest sensitivity to each ROI, the time course of which was extracted and utilised in the subsequent analyses. The procedure was similar to 2.3, except defining CTFs at group-level and extracting ROI timecourses based on subject-level timeseries that were morphed to fsaverage brain (see 2.2) so as to obtain more homogenous evoked responses across subjects for subsequent Fixed Effect Inference on DCMs (see below).

After extraction of the ROI time courses, we used SPM12 (version r6909) for DCM analysis. The model space, as displayed in Figure 3, comprised 28 models. We defined a hierarchical organisation of DCM families in two levels in order to address the following two questions. In the first level of hierarchical comparison, aimed at identification of the winner *grand-family*, all the 28 models were categorised into three families of hub models, no-hub models and no-modulation models. The *hub family* consisted of models 1-16 where one of ATL, IFG, MTG and AG areas played the role of a single hub that received input from the vWFA and was connected to all other semantic areas in the model space. The *no-hub family* consisted of models 17-26. In models 17-18, all the candidate areas where included as multiple convergence zones and received input directly from the vWFA; in models 20-26, only connections of the vWFA to only one of the semantic areas were modulated by the semantic contrast (no further connections to the rest of the network were modulated). Finally, in the *no-modulation* family, models 27-28 had no connection that was modulated by condition (differing only in the presence/absence of self-connections in the vWFA).

**Figure 3.**
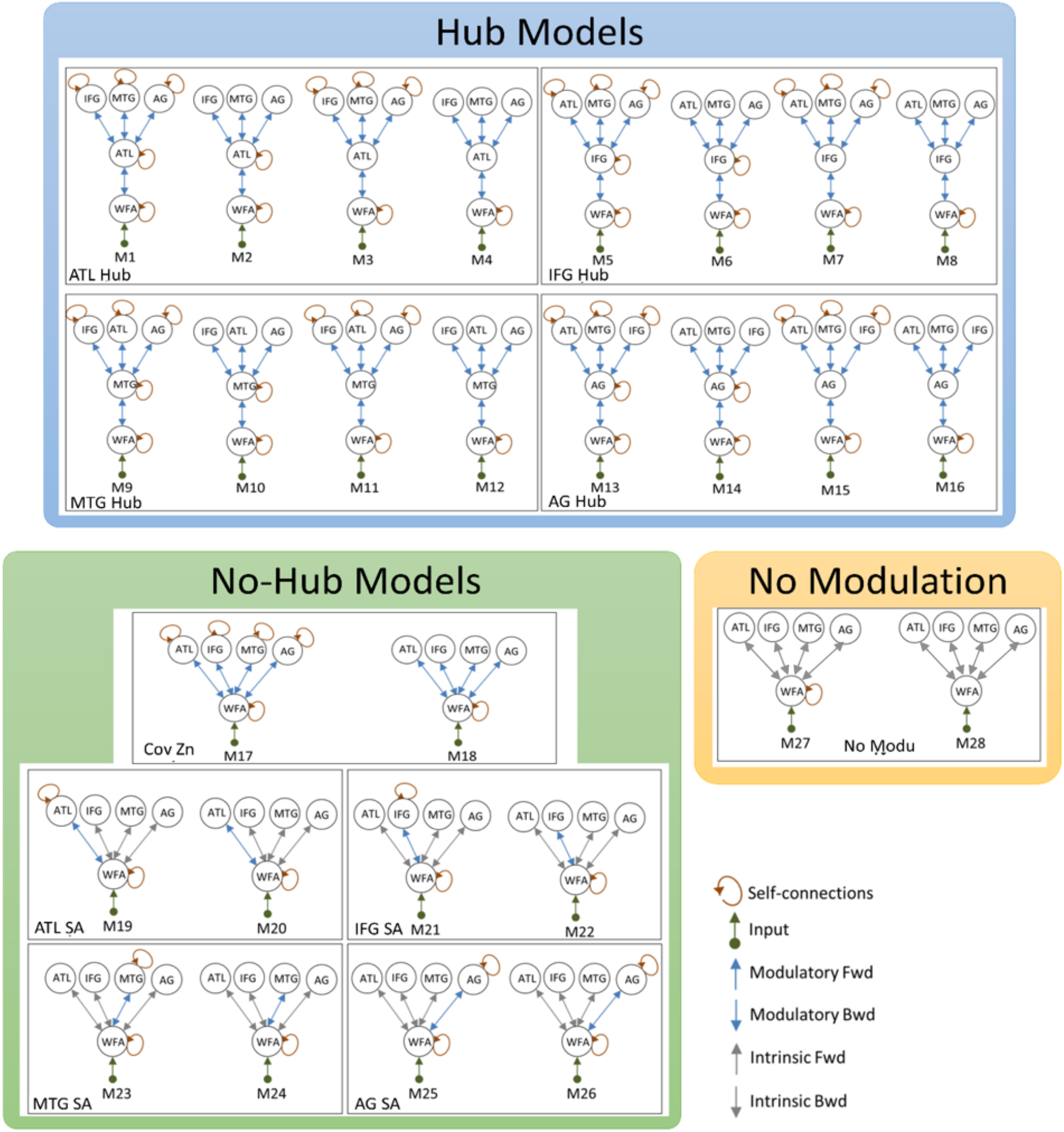
**DCM model space** delineating grand-families and fine-grained families encompassing 28 models. ATL: Anterior Temporal Lobe, IFG: Inferior Frontal Gyrus, MTG: Middle Temporal Gyrus, AG: Angular Gyrus, WFA: Word Form Area, Cov Zn: Convergence Zone, SA: Semantic Area.

Thereafter, in the second level of hierarchical comparison, fine-grained families within the winner grand-family from hierarchy 1 investigations were compared (Figure 3). Models within each fine-grained family spanned different scenarios of self-modulation of the candidate hub areas while self-modulation of vWFA was included in all the models. Finally, we compared single models within the winning fine-grained family in order to examine whether or not one of the models stood out as a conclusive winner and estimated the parameters (i.e. average connection strengths) of this model (details below).

Each model included evoked responses to both concrete and abstract words and was fit for each participant separately. Intrinsic connections were assumed to be common between the conditions, while extrinsic connections were used to model condition-induced modulations of a preselected set of connections. Each model was inverted in two time-windows of 0-250ms and 0-450ms, where the former was considered the early and the latter was considered the later time window. Because DCM is a dynamical system, it requires the data to start at the point of stimulus onset. Thus, both early and late time-windows start from 0ms (i.e. stimulus onset). Data were reduced to 8 spatial modes and no down-sampling, detrending or Hanning windows were used. Furthermore, we used the traditional ERP model for DCM inversion (David *et al.* 2006) instead of the more recently introduced canonical microcircuits (CMC) (Bastos *et al.* 2012) since the former showed higher model evidence for all the models in the model space (see Extended Data Figure 7 - 1). Furthermore, considering the lengths of the time windows of the DCM analysis (i.e. 250ms and 450ms), we included modulations of both forward and backward connections in the model. This choice was made heuristically and informed by the previous literature where semantic effects have been reported as early as 150ms (Moseley *et al.* 2013).

Finally, we used family-level Bayesian Model Selection (BMS) with Fixed Effect Inference (FFX) on the free-energy approximation to the model evidence, in order to identify the winning families in each hierarchy of DCM evaluations (Stephan *et al.* 2007). FFX was considered as more suitable for the current study given that we are studying a homogenous group of healthy young adults, and therefore it is reasonable to assume that the same model applies for all participants. Furthermore, we verified that winning models were not driven by outliers in the free-energy. After conducting BMS, we averaged parameters of the winning single model inside the winning fine-grained family across participants.

## 3 Results

### 3.1 Behavioural results

We conducted paired t-tests in order to compare reaction times (RT) and error rates (ER) for abstract and concrete words. The former showed significantly higher RTs and ERs (RTs for abstract vs concrete: 879±118 vs 778±111ms, ERs: 8.1±5.0 vs 4.5±4.3%), consistent with previous findings (Dhond *et al.* 2007).

### 3.2 Whole-cortex evoked analysis

We used whole-cortex evoked analysis to identify potential representational hubs using a data-driven approach. We defined five time-windows of interest covering the key stages of written word comprehension: 50-150ms, 150-250ms, 250-350ms, 350-450ms, 450-550ms. In the following subsections, we will refer to them by their central time point, e.g. 100ms for 50-150ms.

After source estimation, we examined the grand-average responses to word onset (averaged across concrete and abstract words and across participants), and observed that both conditions showed a posterior-to-anterior flow of current along the ventral occipito-temporal pathway that is typical for visual word recognition (Chen *et al.* 2015; Marinković 2004). Thereafter, using spatio-temporal cluster-based permutations to contrast responses between concrete and abstract words, we found two significant clusters, both including bilateral ATLs and IFGs (Figure 4). The effect started as early as 100ms in the left ATL and extended to the left IFG as well as right ATL/IFG at 300ms, lasting until 450ms. The largest cluster appeared in the time window of 350-450ms. All bilateral clusters showed higher amplitudes for abstract words. Results of the cluster-based permutation and univariate paired t-test are shown in Figure 4.

**Figure 4.**
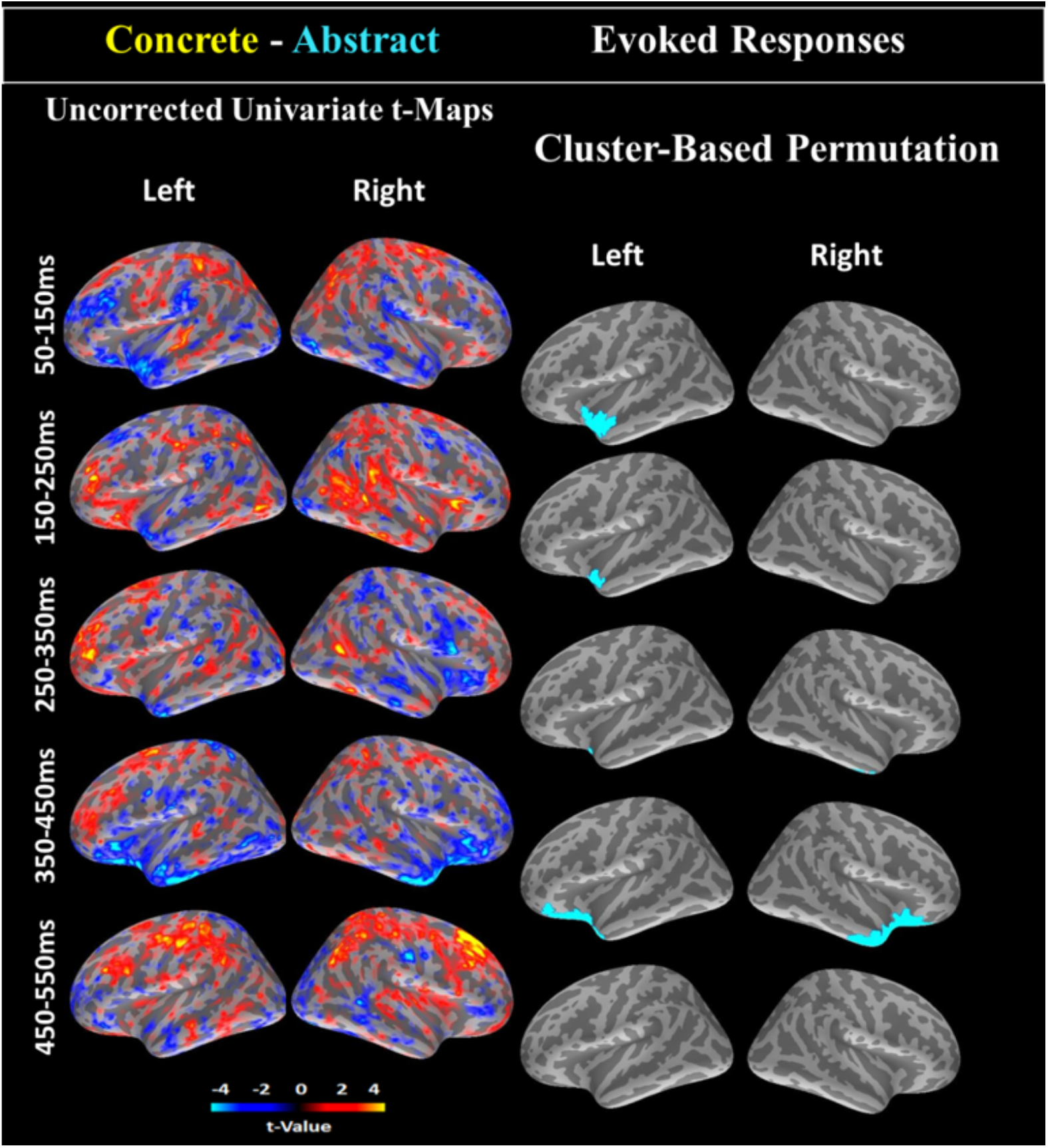
**Whole-cortex evoked responses for concrete minus abstract words:** contrasts of word categories are averaged within five time windows of 100ms duration spanned between 50 and 550ms. Left: uncorrected univariate t-tests. Right: significant clusters of spatio-temporal cluster-based permutation tests. Warm colors indicate higher values for concrete words, and cool colors for the abstract words.

### 3.3 Functional seed-based connectivity

In this analysis we identified the connectivity hubs within the distributed semantic network based on concreteness effects on whole-cortex functional connectivity to sensory-motor-limbic semantic areas. In order to test for these connectivity hubs, we computed seed-based coherence (COH) between the time courses of four candidate semantic hubs in the left ATL, IFG, AG and MTG and every cortical vertex (except those excluded due to EEG/MEG spatial resolution, see Figure 2). Note that a whole-cortex seed-based analysis (i.e. ROI by vertex connectivity) is more informative than an ROI by ROI connectivity approach, considering that concrete and abstract words differ with respect to general but not specific sensory-motor attributes (e.g. concrete words are more tangible but not necessarily action-related), and thus defining sensory-motor ROIs is not straightforward. We calculated connectivity in Theta, Alpha, Beta and Gamma bands, and in two time windows of 150-350ms and 250-450ms. Significant clusters for differentiating concrete and abstract concepts were identified using spatio-temporo-spectral cluster-based permutations. Among the tested seeds, only left ATL and AG showed significant differences in connectivity between concrete and abstract words (Figure 5 a). Left AG showed higher Alpha (250-450ms) and Beta (150-450ms) band COH to the left somatosensory cortex for the concrete words. Left ATL showed higher Beta (250-450ms) and Gamma (150-450ms) band COH to the right orbitofrontal cortex for the abstract words (Figure 5 b). Both effects started in the early time window (150-350ms) and persisted into the later time window (250-450ms).

**Figure 5.**
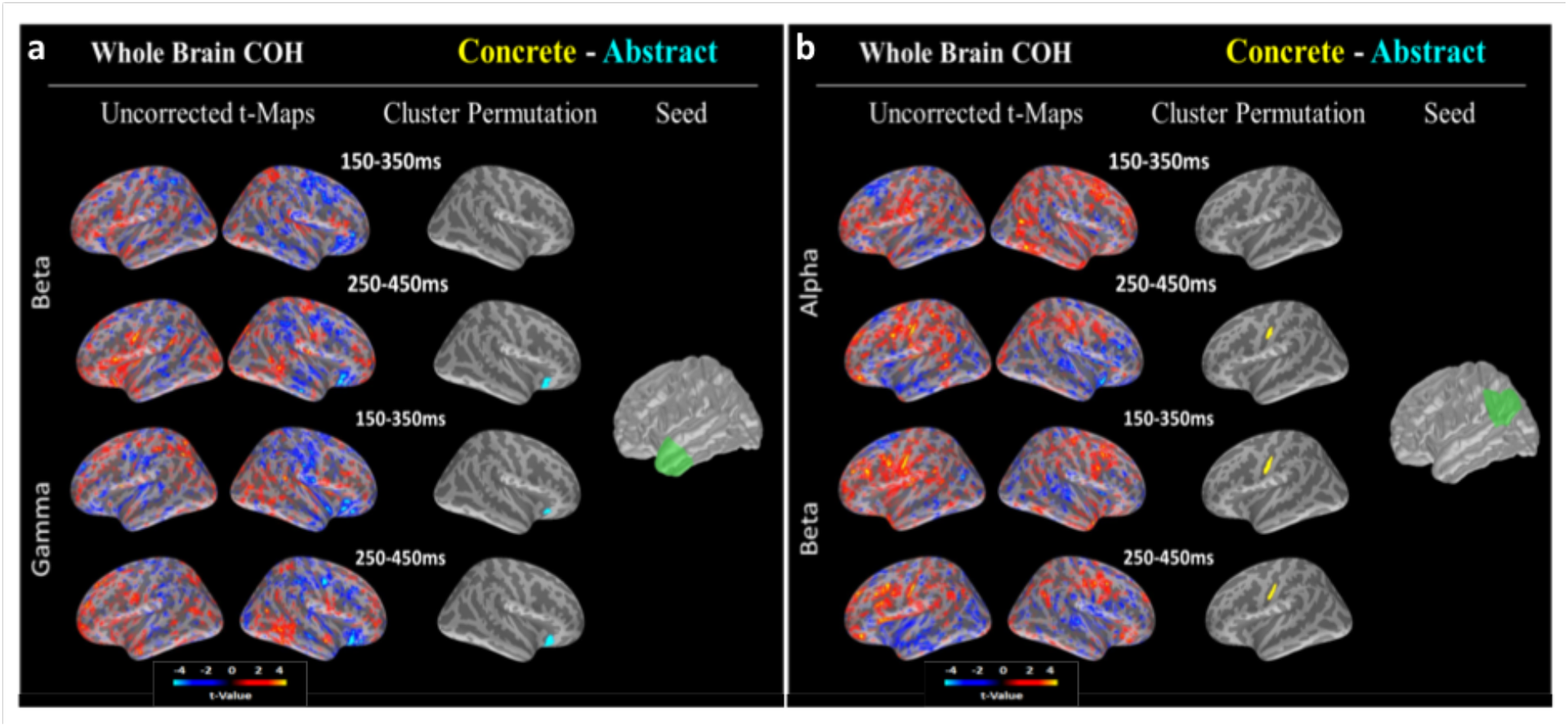
**Results of whole-cortex seed-based connectivity:** a) seed in the left AG. Larger connectivity to left somatosensory cortex for concrete words was found in the beta band for early and late latency ranges, and in the alpha band for the late time window only; b) seed in the left ATL. Larger connectivity to right orbitofrontal cortex for abstract words was found in the gamma band for early and late latency ranges, and in the beta band for the late time window only. Coherence was identified as the suitable connectivity metric for this analysis (see Extended Data Figure 5 - 1 and Table 5 - 1).

### 3.4 Dynamic Causal Modelling

We compared possible organisations of the dynamic network that binds the heteromodal semantic areas using DCM analysis of evoked responses (i.e. ATL, IFG, MTG and AG with vWFA as the input region). The aim was to investigate whether this heteromodal network is bound by a central connectivity hub in the latency ranges 0-250ms and 0-450ms. Heteromodal semantic areas that are involved in semantic processing regardless of the word categories have been established in previous literature. Therefore, we here used DCM, which requires a hypothesis-guided analysis approach (i.e. ROI by ROI connectivity) and can explicitly find the best model of connectivity among the selected areas. As such, hypothesis-guided DCM analysis and data-driven functional connectivity investigations provide complementary evidence to identify the connectivity hubs within the heteromodal semantic network and between the heteromodal nodes and distributed semantic areas, respectively.

#### 3.4.1 ROIs and time courses

ROIs included in the DCM models and their average signed ERP responses for an example participant are shown in Figure 6. We used paired t-tests to test for significant differences between absolute ERP responses to concrete and abstract words in the averaged time windows of exploration (i.e. 0-250ms and 0-450ms) and found that only ATL showed a significant difference between the two conditions (p = 0.016 and p = 0.031 for earlier and later time windows respectively), consistent with the earlier whole-cortex activity analysis.

**Figure 6.**
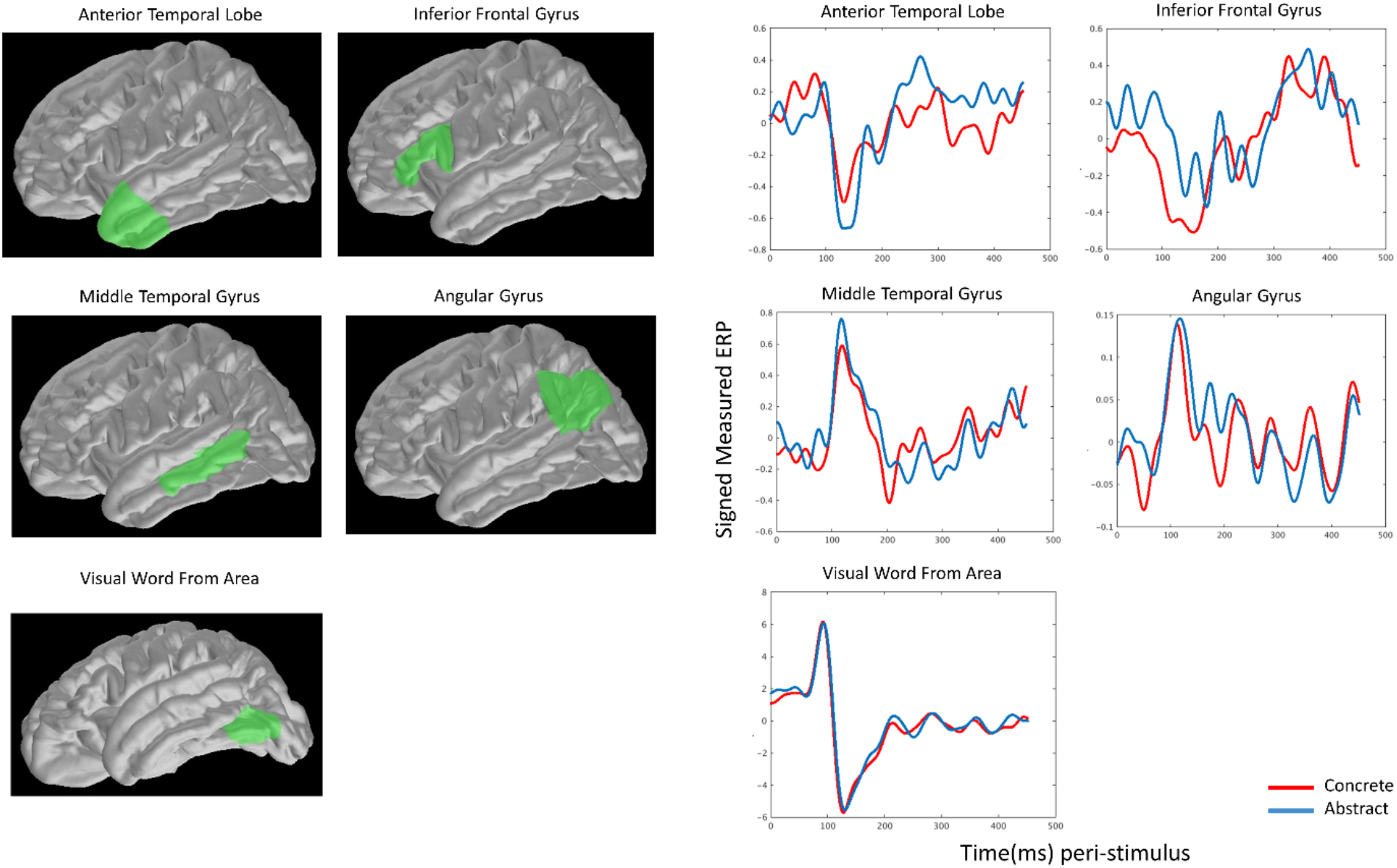
Example signed ERPs for ROIs included in DCM models. Left: ROIs included in the DCM models including ATL, IFG, MTG, AG and vWFA; right: signed ERP time courses of the ROIs from −50ms to 450ms for a representative subject.

#### 3.4.2 First hierarchy: grand-family of hubs showed the highest model evidence

In this step of analysis, we compared three grand-families of models (the hub, no-hub and no-modulation families shown in Figure 3) using FFX BMS and found that the first family showed the highest posterior probability (Figure 7-left panel), within both 0-250ms and 0-450ms post-stimulus windows. This grand-family consisted of four fine-grained families including ATL hub, IFG hub, MTG hub and AG hub. Each of these families consisted of four models where the hub received input from the vWFA and established connections to the other nodes of the heteromodal subnetwork. BMS results are shown in Figure 7-left panel.

**Figure 7.**
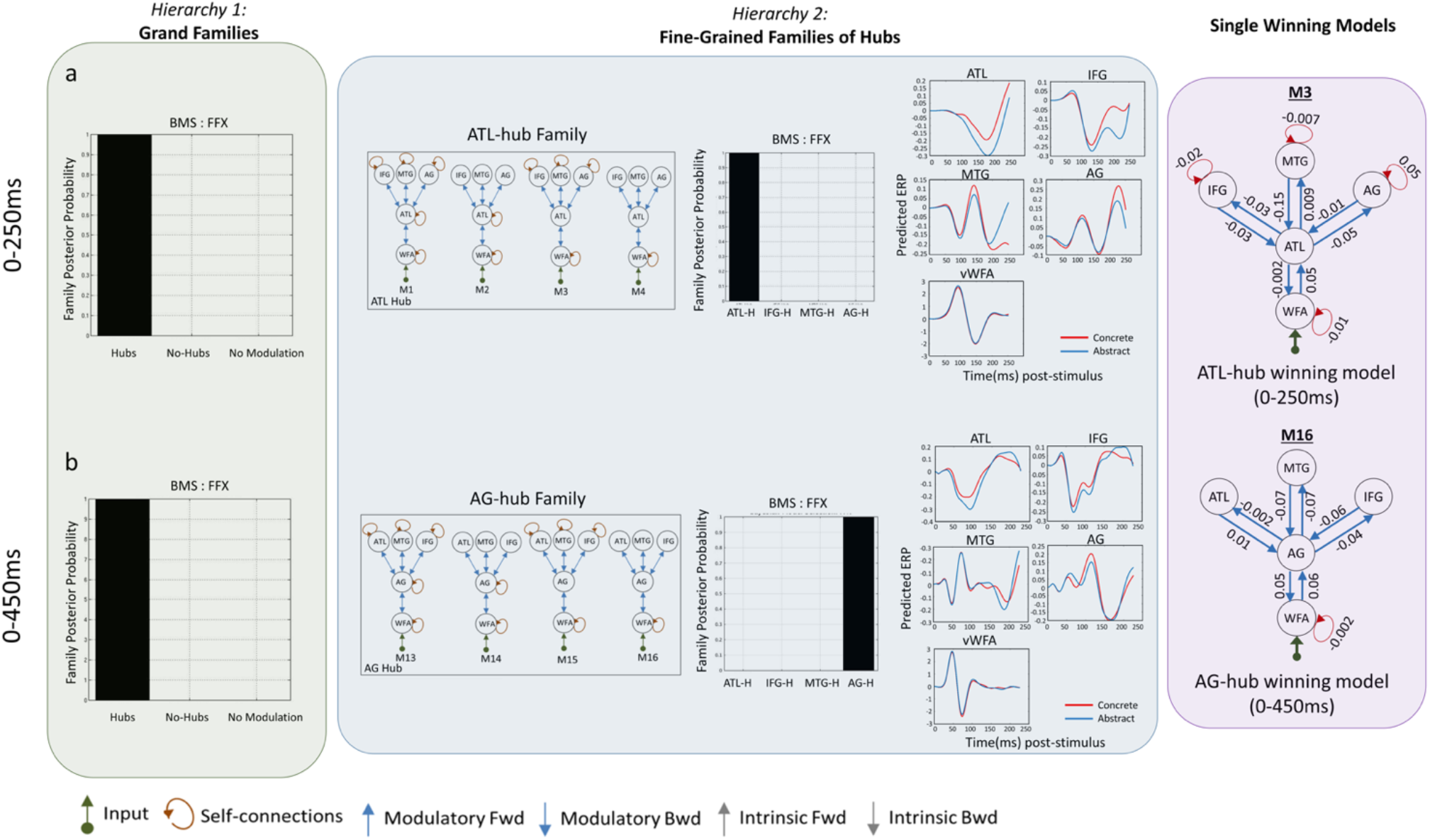
**DCM results based on two hierarchies of family comparisons:** a) DCM results within 0-250ms. Hub models and ATL-hub model were identified as winning families of the first and second hierarchy of comparisons, respectively. Model 3 within the ATL-hub family was identified as a conclusive winner. b) DCM results within 0-450ms. Hub models and AG-hub model were identified as winning families of the first and second hierarchy of comparisons, respectively. Model 4 within the AG-hub family was identified as a conclusive winner. Models were inverted using classic DCM for ERP and not canonical microcircuits (see Extended Data Figure 7 - 1). BMS: Bayesian Model Selection, FFX: Fixed-effect Inference, ATL-H: ATL-Hub, IFG-H: IFG-Hub, MTG-H: MTG-Hub, AG-H: AG-Hub.

#### 3.4.3 Second hierarchy: ATL and AG hub families showed the highest model evidence

In the next step, in order to find areas that serve as a hub, we compared the fine-grained families within the grand-family of hub models, and found that models with ATL as the hub showed the highest posterior probabilities for the 0-250ms time window, and models with AG as the hub showed highest posterior probabilities for the 0-450ms time window (Figure 7-middle panel). In the former family, ATL received input from the vWFA (via bidirectional connections) and was connected to the IFG, MTG and AG. In the latter, AG received input from vWFA and was connected to ATL, IFG and MTG. Each family comprised four models where inter-areal connections were bidirectional and self-modulations of the vWFA were switched on. However, the self-modulation of ATL/AG hubs in their corresponding family, as well as self-modulations of other heteromodal nodes, varied across models in each family (refer to section 2.5 for more details).

##### 3.4.3.1 Single models within ATL and AG hub families

After identifying ATL and AG hubs within 0-250ms and 0-450ms post-stimulus as the winner fine-grained families, we compared models within each family and estimated average parameters across participants for the single winning model within each time-window. Results are shown in Figure 7-right panel. Within the ATL hub family in the 0-250ms time window, model 3 was identified as the conclusive winner (Free energy estimations for the four models: −609.40, −608.12, −526.34, −596.70), where self-connections of vWFA and other areas as well as forward-backward connections of ATL to all the nodes of the network but not self-connections of ATL were modulated. Within AG hub family in 0-450ms time window, model 16 was identified as the conclusive winner (Free energy estimations for the four models: −1751.3, −1941.9, −1738.7, −1578.7). In this model, self-connections of vWFA and forward-backward connections of the AG to all the nodes of the network but not self-connections of AG or convergence zones were allowed to be modulated.

## 4 Discussion

We used source-reconstructed EEG/MEG data in order to characterise spatio-temporal dynamics of semantic brain network using a word concreteness task. Our results provide novel evidence for distinct roles of the anterior temporal lobe (ATL) and angular gyrus (AG) in semantic brain networks, where only ATL appeared as a representational hub, while both ATL and AG appeared as connectivity hubs. Our conclusions are based on three lines of evidence: firstly, our data-driven evoked analysis revealed left ATL to be the first area modulated by word concreteness within 150ms after word presentation. This modulation persisted into later time windows and spread to the bilateral ATLs and inferior frontal gyri (IFGs). We found activation in no other brain areas to be significantly modulated by concreteness, supporting the central role of ATL as a single representational hub. Secondly, whole-cortex seed-based functional connectivity results identified left ATL and AG as connectivity hubs through modulations of connectivity to sensory-motor-limbic systems in two time-windows of 150-350ms and 250-450ms. Thirdly, effective connectivity analysis (dynamic causal modelling, DCM) among the key heteromodal semantic hub candidates (i.e. ATL, IFG, MTG and AG) and with the visual word form area (vWFA) as the input region favoured *single hub* models in both early and later time windows (i.e. 0-250ms and 0-450ms). Comparisons of more fine-grained families of models favoured the left ATL-hub family in the earlier time-window and AG-hub family in the later time-window. Therefore, our results suggest that while both activity and connectivity of ATL are modulated by semantics, especially during earlier stages of semantic information retrieval, AG also supports semantic connectivity at later stages. In the following, we will discuss the implications of our results for the structure of the semantic brain network.

### 4.1 ATL activity is modulated by concreteness

Left ATL’s modulation by concreteness within 150ms post-stimulus provides novel data-driven evidence in support of the key role of this region in semantic processing. This finding is in-line with the hub-and-spokes framework that suggests ATL as a semantic hub that acts as the first link between perceptual and semantic stimulus representations (Lambon Ralph *et al.* 2016; Patterson *et al.* 2007) and thus predicts an early modulation of the hub by semantic variables. Early modulations by lexico-semantic variables have previously been reported in the left ATL and middle temporal gyrus (MTG) (Hauk *et al.* 2012; Westerlund & Pylkkänen 2014) and modality-specific semantic access in sensory-motor cortices (Moseley *et al.* 2013) based on hypothesis-driven ROI-based analyses. However, ROI-based approaches may neglect relevant effects in unexplored regions. Our data-driven whole-cortex approach overcomes these limitations and can be expected to improve the reproducibility and generalisability of our results.

In addition to the earliest modulations of the left ATL, we found the largest effects in the N400 time window (350-450ms) in bilateral ATLs and anterior IFGs. Generally, we found higher absolute activations for abstract words in all clusters/time windows, likely reflecting higher processing demands on the semantic system (Binder *et al.* 2005; Dhond *et al.* 2007; Jackson *et al.* 2015; Lau *et al.* 2013). It is worth noting that ATL activation in this later time window was accompanied by the IFG. While it is possible that our IFG effects are the result of leakage from ATL (Hauk *et al.* 2011; Liu *et al.* 1998), IFG has been implicated in semantics by a range of neuroimaging studies, mostly related to control and unification (Bookheimer 2002; Devlin *et al.* 2003; Lambon Ralph *et al.* 2016). Our abstract words were associated with longer reaction times, which may explain the involvement of cognitive control areas at later stages of processing. Alternatively, the timing of our effects is consistent with previous studies that have suggested a role of IFG in the generation of the N400 ERP component (Hagoort 2004; Lau *et al.* 2008).

### 4.2 Dynamic organisation of the heteromodal semantic subnetwork: key roles of ATL and AG

Prominent models of brain semantic networks propose that a heteromodal subnetwork of semantics is involved in semantic processing through modulation of both activity and connectivity (Binder 2016; Lambon Ralph *et al.* 2016; Pulvermüller 2018; Seghier 2012; Woollams & Patterson 2017) Our results shed new light on dynamic trajectory of connectivity of this network and suggest different types of hubs, namely representational and connectivity hubs within this network.

In our whole-cortex seed-based functional connectivity analyses ATL and AG showed modulations of connectivity from 150 ms post-stimulus. Left ATL was more strongly connected to right orbitofrontal cortex for abstract words, while left AG showed stronger connections to left somatosensory cortex for concrete words. Both regions have been proposed as links to sensory-motor-affective areas, but with a possible advantage of AG for concrete sensorimotor concepts and of ATL for abstract social-affective concepts (Binder *et al.* 2005; Binder & Desai 2011). Interestingly, our abstract words were rated as more emotional (higher valence) than our concrete words, which may have contributed to stronger connections between ATL and orbito-frontal cortex (Binney *et al.* 2016; Lambon Ralph *et al.* 2016). Thus, our observed connectivity patterns are consistent with this proposal.

Additionally, our DCM analysis showed a central role of ATL in an early latency window (0-250 ms), with a conclusive single winning model where all intrinsic and extrinsic connections except for self-connections of the ATL were modulated by the concreteness contrast. Additionally, in the more prolonged time-window (0-450ms), a key role for the AG was revealed, with a conclusive model where the extrinsic but not the intrinsic connections of the semantic areas were modulated. These results suggest that the heteromodal semantic subnetwork is coordinated by a central node that acts as a connector hub bridging between input sensory regions and the rest of semantic network, and suggest that this hub region is dynamically relocated, from ATL to AG, as a concept unfolds in the brain.

### 4.3 Caveats

The choice of connectivity metrics for EEG/MEG analysis is still challenging. Here we used effective and functional connectivity for complementary purposes, namely to identify connectivity hubs within the heteromodal semantic network and between the nodes of this network and all other brain areas, respectively. Coherence was identified as the best metric for functional connectivity in our data based on an empirical approach, but other metrics may give different answers, owing to different sensitivities to different types of neural communication. For effective connectivity, DCM is arguably the sole available method for modelling the full evoked responses to experimental manipulations based on biophysical models of the brain, though even then, there are multiple possible neural models within DCM for EEG/MEG (Bastos *et al.* 2012; David *et al.* 2006). While we used the model evidence to justify the “ERP” model that has been used most commonly in the literature (Chennu *et al.* 2016; David *et al.* 2011; Garrido *et al.* 2008), the model evidence can only find the most likely model among those tested but it cannot determine whether the winning model is in fact the true model. In particular, inspired by the previous literature (Binder *et al.* 2009; Lambon Ralph *et al.* 2016; Martin *et al.* 2014), we here only focused on different scenarios spanning one-layer networks with single or multiple parallel semantic areas or two-layer models with a single hub in the intermediate layer. Nevertheless, current findings do not obviate the possibility of more complex heteromodal semantic networks with different permutations of semantic areas in three or more network layers, potentially involving multiple hubs. Moreover, DCM makes several strong assumptions, the robustness of which needs to be validated in future studies using other datasets, including more direct electrical recordings in humans and animals.

Finally, we utilised a word concreteness decision task in a visual word recognition paradigm which has been successfully and frequently used in previous fMRI (Binder *et al.* 2005) and EEG/MEG (Dhond *et al.* 2007) studies, and can be assumed to involve heteromodal as well as some general modality-specific regions (e.g. somatosensory versus limbic cortices) involved in semantic processing. Some of our findings might be specific to this particular experimental design. Thus, an important next step is to tackle the representational versus connectivity hubs and their timings for more general as well as more specific semantic contrasts, as well as different task settings such as auditory word presentations or nonverbal visual inputs.

### 4.4 Conclusions

Our study provided novel insights into the dynamic brain networks underlying semantic word processing that could not have been provided using metabolic neuroimaging or neuropsychological methods. We confirmed a central role for the ATL as a representational hub that may perform early categorisation and similarity judgments, and for both ATL and AG as connectivity hubs that may support semantic integration and unification processes. Therefore, our results are consistent with both the central role of the ATL as strongly indicated by both patient and imaging work (Lambon Ralph *et al.* 2016; Patterson *et al.* 2007) and also the importance of the AG as indicated by neuroimaging metanalyses (Binder *et al.* 2009). Consideration of the time course of semantic processing in this study has therefore allowed integration of previously distinct approaches to neural semantic networks.

## Acknowledgements

This work was supported by a Cambridge University international scholarship award to S.F and UK Medical Research Council grants to R.N.H. (SUAG/010 RG91365) and O.H. (SUAG/019 RG91365). We would like to thank Dr. Karalyn Patterson for helpful discussions on earlier versions of this work, Dr. Rebecca Jackson for feedback on the manuscript and Drs. Elisa Cooper, Yuanyuan Chen and Gemma Evans for their involvement in data collection.

## 6 Extended Data

**Table 1 - 1.** List of concrete and abstract words used in the experiment.

| Concrete |  |  | Abstract |  |  |
| --- | --- | --- | --- | --- | --- |
| ABDOMEN | JUDGE | TROUT | ACCORD | HAVOC | THRILL |
| ANCHOR | LANTERN | TRUMPET | ACUMEN | HERESY | TOPIC |
| APRICOT | LINEN | VINEGAR | ADAGE | HONESTY | VERITY |
| ARROW | MANTLE | VIOLIN | AMITY | HONOUR | WEALTH |
| BALLOON | MATCH | VOLCANO | APATHY | HORROR | WISDOM |
| BARREL | MEDAL | WHARF | AXIOM | HUMOUR | WRATH |
| BASIN | MUZZLE |  | BELIEF | IDIOM |  |
| BASKET | NAPKIN |  | BOREDOM | IMPETUS |  |
| BAYONET | NECTAR |  | BRAVERY | INSIGHT |  |
| BLANKET | NOZZLE |  | CHEAT | INTERIM |  |
| BLISTER | NUTMEG |  | COURAGE | LEVITY |  |
| BLOSSOM | OLIVE |  | CUSTOM | LIBERTY |  |
| CABINET | ONION |  | DEITY | LOYALTY |  |
| CAMERA | ORANGE |  | DELIGHT | MALADY |  |
| CELLAR | OYSTER |  | DEMON | MASTERY |  |
| CHAPEL | PEDAL |  | DENIAL | MERIT |  |
| CHISEL | PERCH |  | DESPAIR | MIRACLE |  |
| CHROME | PIANO |  | DIGNITY | MISERY |  |
| CLOVE | PIGEON |  | DOGMA | MOTIVE |  |
| COWHIDE | PLANK |  | DREAM | MYSTERY |  |
| CRADLE | PLATTER |  | EMOTION | NADIR |  |
| CROSS | PORCH |  | ENIGMA | PANIC |  |
| DIAMOND | PRISON |  | FALLACY | PARADOX |  |
| DOUGH | PRUNE |  | FANTASY | PASSION |  |
| EAGLE | PULPIT |  | FETISH | PAUSE |  |
| EASEL | RIFLE |  | FLAIR | PHANTOM |  |
| EMERALD | ROCKET |  | FOLLY | PIQUE |  |
| FLASK | RUDDER |  | FORFEIT | RARITY |  |
| FLOWER | SADDLE |  | FORTUNE | REALM |  |
| FOREST | SATIN |  | GAIETY | REGRET |  |
| FRAME | SEAWEED |  | GALLANT | REVENGE |  |
| FURNACE | SHAWL |  | GENIUS | ROMANCE |  |
| GALLERY | SHUTTER |  | GLOOM | SCARE |  |
| GIRDLE | SKYLARK |  | GLORY | SCORN |  |
| GLACIER | SNAKE |  | GRACE | SHAME |  |
| GRAVEL | STADIUM |  | GREED | SIMILE |  |
| HAMMER | STEEPLE |  | GRIEF | SORROW |  |
| HATCHET | STRAW |  | GRUDGE | SPITE |  |
| ICICLE | SWORD |  | GUILT | SPREE |  |
| IODINE | TEMPLE |  | HARMONY | TEMPER |  |
| IVORY | TRIPOD |  | HATRED | THEME |  |

**Table 5 - 1.** PCA analysis of seed-based connectivity across connectivity methods. Based on these results, Coherence was identified as the most suitable method for functional connectivity analysis in Figure 5.

| Pooled<br>Con/Abs |  |  |  |  |  |  |  |  |  |  |  |  |
| --- | --- | --- | --- | --- | --- | --- | --- | --- | --- | --- | --- | --- |
| PCA across connectivity methods |  |  |  |  |  |  |  |  |  |  |  |  |
|  | 1<sup>st</sup> PC |  |  |  | 2<sup>nd</sup> PC |  |  |  | 3<sup>rd</sup> PC |  |  |  |
|  | Theta | Alpha | Beta | Gamma | Theta | Alpha | Beta | Gamma | Theta | Alpha | Beta | Gamma |
| %EV | 54.7±3.8 | 55.9±3.7 | 61.1±4.5 | 62.1±4.6 | 31±1.5 | 30.1±1.7 | 28.2±3.4 | 26.3±3.3 | 14.2±2.9 | 14±3 | 10.8±2.7 | 11.6±2.7 |
| MI | 0.37±0.1 | 0.44±0.1 | 0.5±0.2 | 0.58±0.1 | -0.87±0.3 | -0.89±0.05 | 0.82±0.2 | 0.8±0.1 | 0.003±0.04 | 0.003±0.03 | 0.01±0.07 | 0.02±0.06 |
| PPC | 0.86±0.03 | 0.86±0.03 | 0.86±0.1 | 0.87±0.03 | 0.2±0.09 | 0.23±0.06 | -0.24±0.2 | -0.27±0.1 | 0.46±0.05 | 0.45±0.05 | 0.23±0.3 | 0.29±0.3 |
| COH | 0.86±0.03 | 0.86±0.03 | 0.87±0.2 | 0.87±0.04 | 0.19±0.06 | 0.22±0.05 | -0.21±0.1 | -0.26±0.1 | -0.46±0.05 | -0.46±0.05 | -0.23±0.3 | -0.3±0.3 |

EV: Explained Variance; MI: Mutual Information; PPC: Pairwise Phase Consistency; COH: Coherence

**Figure 5 - 1.**
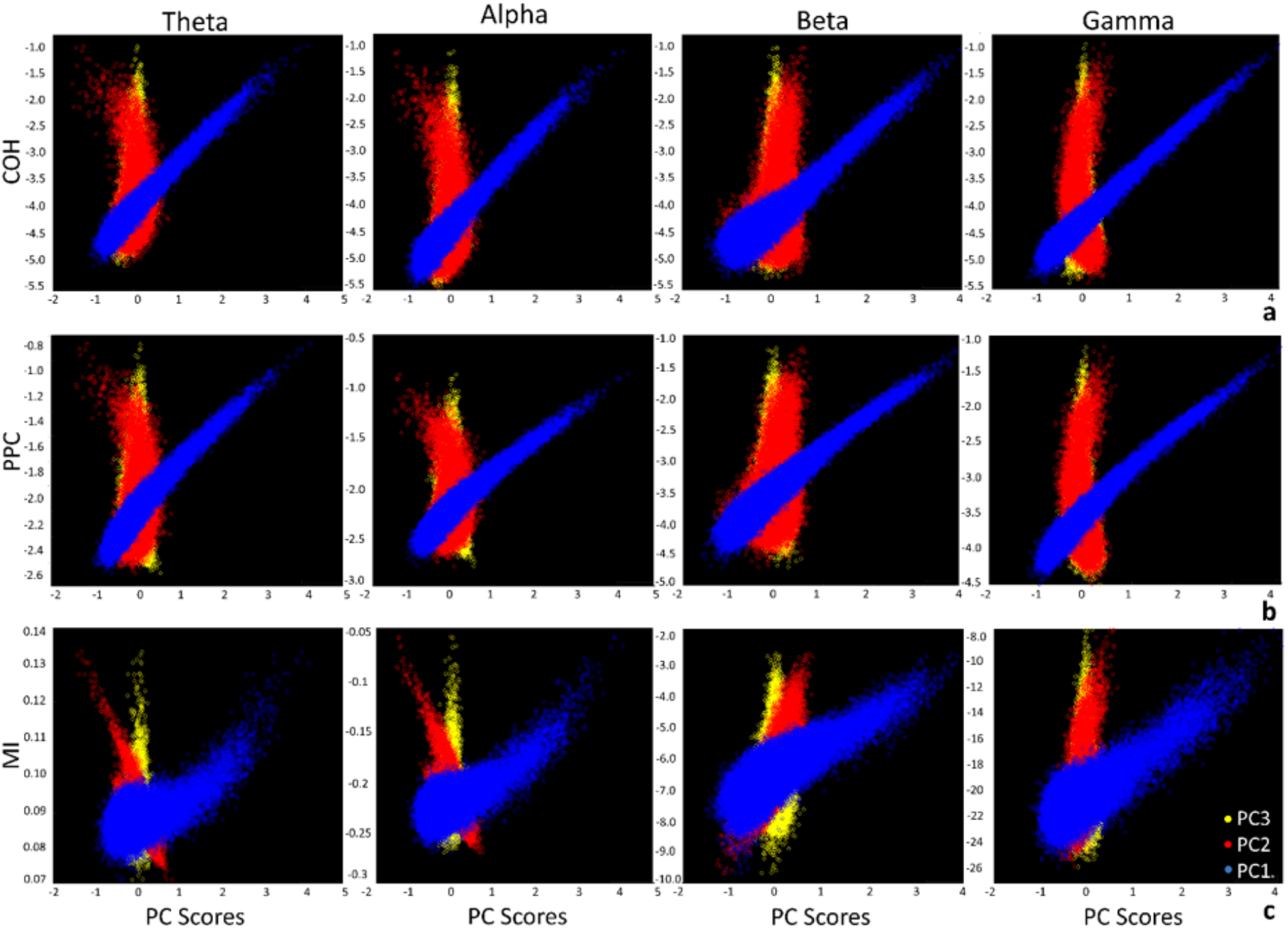
PCA analysis of seed-based connectivity across connectivity methods. Boxcox-transformed a) COH; b) PPC and c) MI plotted against distributions of the first (blue), second (red) and third (yellow) principal component scores over vertices and time windows (averaged across seeds and participants). The first PC showed high linear correlation with COH and PPC (>0.8) while the second PC showed higher correlation with MI. More details are presented in Table 5 - 1. Based on these results, Coherence was identified as the most suitable method for functional connectivity analysis in Figure 5. COH: Coherence, PPC: Pairwise Phase Consistency, MI: Mutual Information, PCA: Principal Component Analysis.

**Figure 7 - 1.**
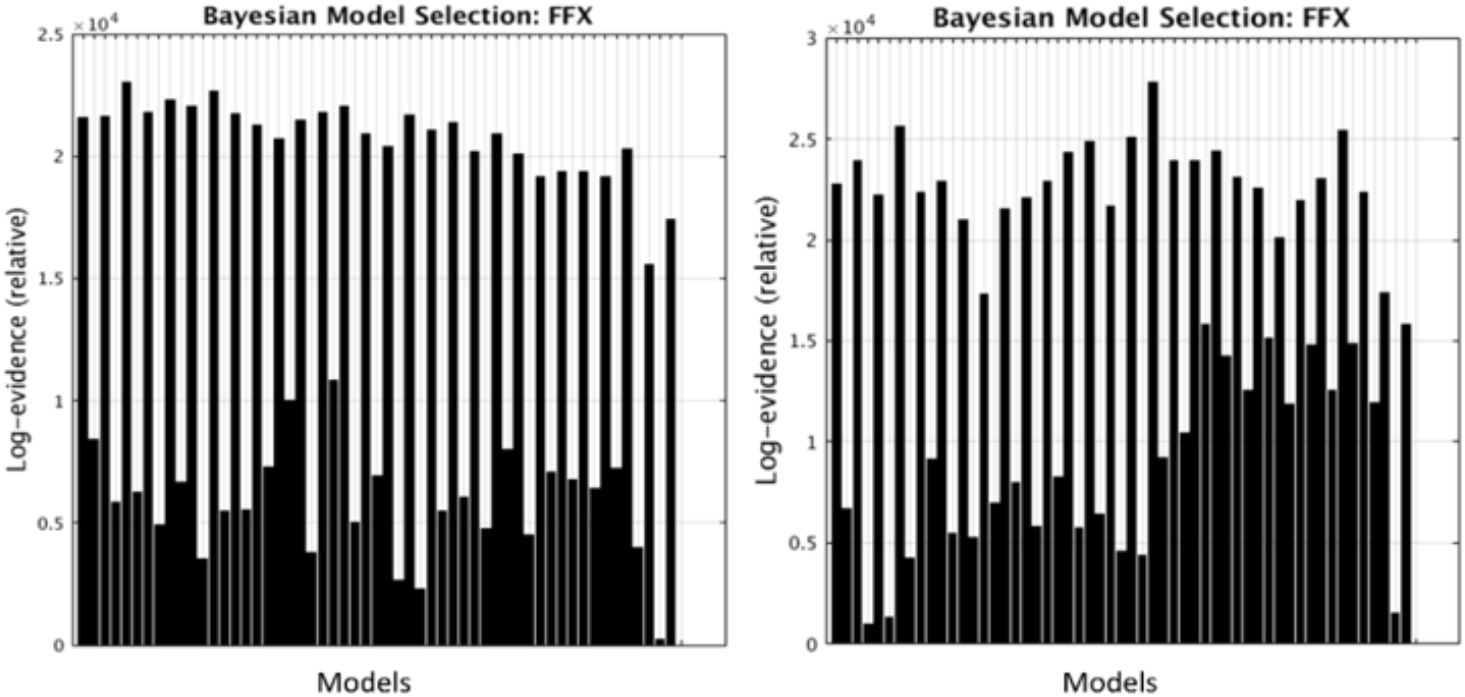
Comparing model evidence of classic DCM for ERP (David *et al.* 2006) and canonical microcircuit models (CMC) (Bastos *et al.* 2012) for the 28 models in Figure 3 within 0-250ms (left) and 0-450ms (right). Results of analysis are shown in Figure 7. Odd numbers show ERP results while even numbers show CMC results. Therefore, the former shows substantially higher model evidence.

## Bibliography

Baker, A. P., Brookes, M. J., Rezek, I. a., … Woolrich, M. (2014). Fast transient networks in spontaneous human brain activity. ELife, 2014 (3), 1–18.

Barsalou, L. W. (2009). Simulation, situated conceptualization, and prediction. Philosophical Transactions of the Royal Society B: Biological Sciences, 364(1521), 1281–1289.

Bastos, A. M., & Schoffelen, J.-M. (2016). A Tutorial Review of Functional Connectivity Analysis Methods and Their Interpretational Pitfalls. Frontiers in Systems Neuroscience, 9(January), 1–23.

Bastos, A. M., Usrey, W. M., Adams, R. a., Mangun, G. R., Fries, P., & Friston, K. J. (2012). Canonical Microcircuits for Predictive Coding. Neuron, 76(4), 695–711.

Binder, J. R. (2016). In defense of abstract conceptual representations. Psychonomic Bulletin & Review, 23, 1096–1108.

Binder, J. R., & Desai, R. H. (2011). The neurobiology of semantic memory. Trends in Cognitive Sciences, 15(11), 527–36.

Binder, J. R., Desai, R. H., Graves, W. W., & Conant, L. L. (2009). Where is the semantic system? A critical review and meta-analysis of 120 functional neuroimaging studies. Cerebral Cortex (New York, N.Y.: 1991), 19(12), 2767–96.

Binder, J. R., Westbury, C. F., Mckiernan, K. A., Possing, E. T., & Medler, D. A. (2005). Distinct brain systems for processing concrete and abstract concepts. Journal of cognitive neuroscience, 17(6), 905–917.

Binney, R. J., Hoffman, P., & Lambon Ralph, M.a. (2016). Mapping the Multiple Graded Contributions of the Anterior Temporal Lobe Representational Hub to Abstract and Social Concepts: Evidence from Distortion-corrected fMRI. Cerebral Cortex, 26(11), 4227–4241.

Bookheimer, S. (2002). Functional MRI of Language: New Approaches to Understanding the Cortical Organization of Semantic Processing. Annual Review of Neuroscience, 25(1), 151–188.

Brookes, M. J., Woolrich, M., Luckhoo, H., … Morris, P. G. (2011). Investigating the electrophysiological basis of resting state networks using magnetoencephalography. Proceedings of the National Academy of Sciences of the United States of America, 108(40), 16783–8.

Bullmore, E., & Sporns, O. (2009). Complex brain networks: graph theoretical analysis of structural and functional systems. Nature Reviews Neuroscience, 10(3), 186–98.

Chen, Y., Davis, M. H., Pulvermüller, F., & Hauk, O. (2015). Early Visual Word Processing Is Flexible: Evidence from Spatiotemporal Brain Dynamics. Journal of Cognitive Neuroscience, 27(9), 1738– 1751.

Chennu, S., Noreika, V., Gueorguiev, D., Shtyrov, Y., Bekinschtein, T. A., & Henson, R. (2016). Silent Expectations: Dynamic Causal Modeling of Cortical Prediction and Attention to Sounds That Weren’t. Journal of Neuroscience, 36(32), 8305–8316.

Colclough, G. L., Brookes, M. J., Smith, S. M., & Woolrich, M. W. (2015). A symmetric multivariate leakage correction for MEG connectomes. NeuroImage, 117, 439–448.

Coltheart, M. (1981). The mrc psycholinguistic database. The Quarterly Journal of Experimental Psychology Section A, 33(4), 497–505.

Dale, a M., Fischl, B., & Sereno, M. I. (1999). Cortical surface-based analysis. I. Segmentation and surface reconstruction. NeuroImage, 9(2), 179–194.

Daub, C. O., Steuer, R., Selbig, J., … Stoughton, R. (2004). Estimating mutual information using B-spline functions – an improved similarity measure for analysing gene expression data. BMC Bioinformatics, 5(1), 118.

David, O., Kiebel, S. J., Harrison, L. M., Mattout, J., Kilner, J. M., & Friston, K. J. (2006). Dynamic causal modeling of evoked responses in EEG and MEG. NeuroImage, 30(4), 1255–1272.

David, O., Maess, B., Eckstein, K., & Friederici, A. D. (2011). Dynamic causal modeling of subcortical connectivity of language. The Journal of Neuroscience, 31(7), 2712–2717.

Devlin, J. T., Matthews, P. M., & Rushworth, M. F. S. (2003). Semantic Processing in the Left Inferior Prefrontal Cortex: A Combined Functional Magnetic Resonance Imaging and Transcranial Magnetic Stimulation Study. Journal of Cognitive Neuroscience, 15(1), 71–84.

Dhond, R. P., Witzel, T., Dale, A. M., & Halgren, E. (2007). Spatiotemporal cortical dynamics underlying abstract and concrete word reading. Human Brain Mapping, 28(4), 355–62.

Farahibozorg, S. R., Henson, R. N., & Hauk, O. (2018). Adaptive cortical parcellations for source reconstructed EEG/MEG connectomes. NeuroImage, 169, 23–45.

Fischl, B., Sereno, M. I., & Dale, A. M. (1999). Cortical Surface-Based Analysis II: Inflation, Flattening, and a Surface-Based Coordinate System. NeuroImage, 9(2), 195–207.

Garrido, M. I., Friston, K. J., Kiebel, S. J., Stephan, K. E., Baldeweg, T., & Kilner, J. M. (2008). The functional anatomy of the MMN: A DCM study of the roving paradigm. NeuroImage, 42(2), 936–944.

Gramfort, A., Luessi, M., Larson, E., … Hämäläinen, M. (2013). MEG and EEG data analysis with MNE-Python. Frontiers in Neuroscience, 7, 267.

Gramfort, A., Luessi, M., Larson, E., … Hämäläinen, M. S. (2014). MNE software for processing MEG and EEG data. NeuroImage, 86(2014), 446–460.

Greenblatt, R. E., Pflieger, M. E., & Ossadtchi, A. E. (2012). Connectivity measures applied to human brain electrophysiological data. Journal of Neuroscience Methods, 207(1), 1–16.

Hagoort, P., Hald, L., Bastiaansen, M., & Petersson, K. M. (2004). Integration of word meaning and world knowledge in language comprehension. Science, 304(5669), 438–441.

Hauk, O. (2016). Only time will tell – why temporal information is essential for our neuroscientific understanding of semantics. Psychonomic Bulletin and Review, 23(4), 1072–1079.

Hauk, O., Coutout, C., Holden, A., & Chen, Y. (2012). The time-course of single-word reading: Evidence from fast behavioral and brain responses. NeuroImage, 60(2), 1462–1477.

Hauk, O., Wakeman, D. G., & Henson, R. (2011). Comparison of noise-normalized minimum norm estimates for MEG analysis using multiple resolution metrics. NeuroImage, 54(3), 1966–1974.

Hyvärinen, A., & Oja, E. (2000). Independent component analysis: algorithms and applications. Neural Networks, 13(4–5), 411–30.

Jackson, R. L., Bajada, C. J., Rice, G. E., Cloutman, L. L., & Lambon Ralph, M.A. (2018). An emergent functional parcellation of the temporal cortex. NeuroImage, 170, 385–399.

Jackson, R. L., Lambon Ralph, M.A., & Pobric, G. (2015). The Timing of Anterior Temporal Lobe Involvement in Semantic Processing. Journal of Cognitive Neuroscience, 27(7), 1388–1396.

Jung, T. P., Makeig, S., Humphries, C., … Sejnowski, T. J. (2000). Removing electroencephalographic artifacts by blind source separation. Psychophysiology, 37(2), 163–178.

Lachaux, J., Rodriguez, E., Martinerie, J., & Varela, F. J. (1999). Measuring Phase Synchrony in Brain Signals. Human brain mapping, 8(4), 194–208.

Lagerlund, T. D., Sharbrough, F. W., & Busacker, N. E. (1997). Spatial filtering of multichannel electroencephalographic recordings through principal component analysis by singular value decomposition. Journal of clinical neurophysiology, 14(1), 73–82.

Lambon Ralph, M.a, Jefferies, E., Patterson, K., & Rogers, T. T. (2016). The neural and computational bases of semantic cognition. Nature Reviews Neuroscience, 18(1), 1–14.

Lau, E. F., Gramfort, A., Hämäläinen, M. S., & Kuperberg, G. R. (2013). Automatic semantic facilitation in anterior temporal cortex revealed through multimodal neuroimaging. The Journal of Neuroscience, 33(43), 17174–81.

Lau, E. F., Phillips, C., & Poeppel, D. (2008). A cortical network for semantics: (de)constructing the N400. Nature Reviews Neuroscience, 9(12), 920–933.

Liu, A. K., Belliveau, J. W., & Dale, A. M. (1998). Spatiotemporal imaging of human brain activity using functional MRI constrained magnetoencephalography data: Monte Carlo simulations. Proceedings of the National Academy of Sciences of the United States of America, 95(15), 8945–8950.

Marinković, K. (2004). Spatiotemporal dynamics of word processing in the human cortex. The Neuroscientist, 10(2), 142–52.

Maris, E., & Oostenveld, R. (2007). Nonparametric statistical testing of EEG- and MEG-data. Journal of Neuroscience Methods, 164(1), 177–90.

Martin, A. (2016). GRAPES—Grounding representations in action, perception, and emotion systems: How object properties and categories are represented in the human brain. Psychonomic Bulletin and Review, 23(4), 979–990.

Martin, A., Kyle Simmons, W., Beauchamp, M. S., & Gotts, S. J. (2014). Is a single “hub”, with lots of spokes, an accurate description of the neural architecture of action semantics?: Comment on ‘Action semantics: A unifying conceptual framework for the selective use of multimodal and modality-specific object knowledge’. Physics of Life Reviews, 11(2), 261–2.

Meyer, K., & Damasio, A. (2009). Convergence and divergence in a neural architecture for recognition and memory. Trends in Neurosciences, 32(7), 376–382.

Molins, A., Stufflebeam, S. M., Brown, E. N., & Hämäläinen, M. S. (2008). Quantification of the benefit from integrating MEG and EEG data in minimum ℓ2-norm estimation. NeuroImage, 42(3), 1069–1077.

Moseley, R. L., Pulvermüller, F., & Shtyrov, Y. (2013). Sensorimotor semantics on the spot: brain activity dissociates between conceptual categories within 150 ms. Scientific Reports, 3, 1928.

Nunez, P. L., Srinivasan, R., Westdorp, A. F., … Cadusch, P. J. (1997). EEG coherency. Electroencephalography and Clinical Neurophysiology, 103(5), 499–515.

Patterson, K., Nestor, P. J., & Rogers, T. T. (2007). Where do you know what you know? The representation of semantic knowledge in the human brain. Nature Reviews Neuroscience, 8(12), 976–87.

Pedregosa, F., Varoquaux, G., Gramfort, A., … Duchesnay, É. (2011). Scikit-learn: Machine Learning in Python. Journal of Machine Learning Research, 12(Oct), 2825–2830.

Phillips, H. N., Blenkmann, A., Hughes, L. E., Bekinschtein, T. A., & Rowe, J. B. (2015). Hierarchical Organization of Frontotemporal Networks for the Prediction of Stimuli across Multiple Dimensions. Journal of Neuroscience, 35(25), 9255–9264.

Pulvermüller, F. (2013). How neurons make meaning: brain mechanisms for embodied and abstract-symbolic semantics. Trends in Cognitive Sciences, 17(9), 458–70.

Pulvermüller, F. (2018). Neural reuse of action perception circuits for language, concepts and communication. Progress in Neurobiology, 160, 1–44.

Pulvermüller, F., Shtyrov, Y., & Hauk, O. (2009). Understanding in an instant: Neurophysiological evidence for mechanistic language circuits in the brain. Brain and Language, 100(2), 81–94.

Rice, G. E., Ralph, M. A. L. L., & Hoffman, P. (2015). The roles of left versus right anterior temporal lobes in conceptual knowledge: An ALE meta-analysis of 97 functional neuroimaging studies. Cerebral Cortex, 25(11), 4374–4391.

Rogers, T. T., Lambon Ralph, M., Garrard, P., … Patterson, K. (2004). Structure and deterioration of semantic memory: a neuropsychological and computational investigation. Psychological Review, 111(1), 205–35.

Sakia, R. M. (1992). The Box-Cox Transformation Technique: A Review. The Statistician, 41(2), 169.

Samu Taulu and Matti Kajola. (2005). Presentation of electromagnetic multichannel data: The signal space separation method. Journal of Applied Physics, 97(12).

Seghier, M. (2012). The Angular Gyrus: Multiple Functions and Multiple Subdivisions. The Neuroscientist, 19(1), 43–61.

Segonne, F., Dale, a M., Busa, E., … Fischl, B. (2004). A hybrid approach to the skill stripping problem in MRI. NeuroImage, 22, 1060–1075.

Sharon, D., Hämäläinen, M. S., Tootell, R. B. H., Halgren, E., & Belliveau, J. W. (2007). The advantage of combining MEG and EEG: Comparison to fMRI in focally stimulated visual cortex. NeuroImage, 36(4), 1225–1235.

Smith, S. M., & Nichols, T. E. (2009). Threshold-free cluster enhancement: Addressing problems of smoothing, threshold dependence and localisation in cluster inference. NeuroImage, 44(1), 83–98.

Snowden, J. S., Harris, J. M., Thompson, J. C., … Neary, D. (2017). Semantic dementia and the left and right temporal lobes. Cortex, 1–16.

Stephan, K. E., Weiskopf, N., Drysdale, P. M., Robinson, P. A., & Friston, K. J. (2007). Comparing hemodynamic models with DCM. NeuroImage, 38(3), 387–401.

Tomasello, R., Garagnani, M., Wennekers, T., & Pulvermüller, F. (2017). Brain connections of words, perceptions and actions: A neurobiological model of spatio-temporal semantic activation in the human cortex. Neuropsychologia, 98, 111–129.

Vinck, M., van Wingerden, M., Womelsdorf, T., Fries, P., & Pennartz, C. M. A. (2010). The pairwise phase consistency: A bias-free measure of rhythmic neuronal synchronization. NeuroImage, 51(1), 112–122.

Westerlund, M., & Pylkkänen, L. (2014). The role of the left anterior temporal lobe in semantic composition vs. semantic memory. Neuropsychologia, 57(1), 59–70.

Woollams, A. M., & Patterson, K. (2017). Cognitive consequences of the left-right asymmetry of atrophy in semantic dementia. Cortex, 107(2018): 64–77.

